# Establishing a straightforward I-SceI mediated recombination one plasmid system for efficient genome editing in *P. putida* KT2440

**DOI:** 10.1101/2024.01.23.576838

**Authors:** Hao Meng, Sebastian Köbbing, Lars M. Blank

## Abstract

*Pseudomonas putida* has become an increasingly important chassis for the production of valuable bioproducts. This development is not at least due to the ever-improving genetic toolbox, including gene and genome editing techniques. Here, we present a novel, one plasmid design of a key genetic tool, the pEMG/pSW system, guaranteeing one engineering cycle to be finalized in three days. The pEMG/pSW system proved in the last decade to be valuable for targeted gensome engineering in *Pseudomonas*, as it enables the deletion of large regions of the genome, the integration of heterologous gene clusters or targeted generation of point mutations. Here, to expedite genetic engineering, two alternative plasmids were constructed: 1) the *sacB* gene from *Bacillus subtilis* was integrated into the I-SceI expressing plasmid pSW-2 as counterselection marker to accelerated plasmid curing; 2) double strand break introducing gene I-SceI and SacB counterselection marker were integrated into the backbone of the original pEMG vector, named pEMG-RIS. The single plasmid of pEMG-RIS allows rapid genome editing despite the low transcriptional activity of a single copy of the I-SceI encoding gene. Here, the usability of the pEMG-RIS is shown in *P. putida* KT2440 by integrating an expression cassette including a *msfGFP* gene in three days. In addition, a large fragment of almost 16 kb was also integrated. In summary, an updated pEMG/pSW genome editing system is presented that allows efficient and rapid genome editing in *P. putida*. The pEMG-RIS will be available via the Addgene platform.

## Introduction

*Pseudomonas putida*, a gram-negative soil bacterium characterized by its versatile metabolism and remarkable stress resistance, has become an increasingly important chassis (Martínez-García, *et al*., 2014, Nikel and de Lorenzo, 2014, Nikel and de Lorenzo, 2018, Weimer, *et al*., 2020) for the production of value-added products such as aromatics (Schwanemann, *et al*., 2020) and glycolipids (Tiso, *et al*., 2020) from renewable carbon source or waste streams (Ballerstedt, *et al*., 2021). In recent years, several markerless genome editing methods have been developed and are extensively applied in the metabolic and genetic engineering of *Pseudomonas* (Galvão and de Lorenzo, 2005, Martínez-García and de Lorenzo, 2011, Luo, *et al*., 2016, Aparicio, *et al*., 2019). One prominent tool is homologous recombination-based editing tools, like the pEMG/pSW system established by Martínez-García *et al*. The system is very well suited for successive scarless deletions, insertions or point mutations in *Pseudomonas* (Martínez-García and de Lorenzo, 2011, Wirth, *et al*., 2020). The method based on two plasmids: a suicide vector with two I-SceI sites and a second vector providing the I-SceI endonuclease. Briefly, the utilization of this system for genome editing requires integration of the pEMG vector into the genome by homologous recombination. An I-SceI encoding plasmid transferred to the co-integrate strain and mediating double strand breaks (DSB) in the I-SceI sites. The DSB are repaired by homologous recombination and the co-integrate is flipped out. The entire process typically takes approximately 6 days by using triparental mating plus some additional days for plasmid curing after confirmation of correct mutants (Volke, *et al*., 2020, Wirth, *et al*., 2020, Volke, *et al*., 2021). It appears that there is potential for further optimization. Recently, cytidine deaminase-based toolsets that enable efficient multiplex editing in *P. putida* have been developed (Volke, *et al*., 2022, Yue, *et al*., 2022, Kozaeva, *et al*., 2024). However, this kind of approach necessitates several prerequisites and has the potential to introduce unknown mutations beyond the spacer region (Volke, *et al*., 2022).

The implementation of counterselection strategies is frequently contemplated and utilized in *Pseudomonas*, such as *pyrF/URA3* (Galvão and de Lorenzo, 2005), UPRTase (Graf and Altenbuchner, 2011), and Cre*/loxP* (Luo, *et al*., 2016). Reyrat *et al*. presented an extensive overview of diverse counterselection markers, including *sacB*, *ccdB*, and *pheS* (Reyrat, *et al*., 1998). The *sacB* gene from *Bacillus subtilis*, which encodes the enzyme levansucrase responsible for the conversion of sucrose into levans, is frequently employed as a counterselection marker. When strains harboring the *sacB* gene are plated in the presence of sucrose, the accumulation of levans in the periplasm of gram-negative bacteria can cause death (Steinmetz, *et al*., 1983).

In this study, we demonstrate the construction and applicability of a one plasmid system for efficient genome editing in *P. putida* KT2440, based on the widely used pEMG/pSW system. The system comprises the pEMG vector backbone, an I-SceI endonuclease regulated by a stringent inducible promoter, and *sacB* gene as a counterselection marker. The new plasmid is integrated into the genome by homologous recombination and I-SceI expression can be precisely induced, without the need of the second plasmid delivery. Additionally, sucrose can facilitate the selection for clones with successfully released co-integrates from the genome. The streamlined method enables the introduction of genomic modification in *P. putida* within 3 days and obtains plasmid-free mutants. To the best of our knowledge, this study presents the most rapid genome editing method in *P. putida* KT2440 reported thus far.

## Experimental procedures

### Bacterial strains, chemicals, and DNA manipulations

The plasmids and bacterial strains used and generated in this study are listed in Table 1 and Table 2. All oligonucleotides used in this study are listed in Table S1. *E. coli* DH5α λ*pir* was used as a cloning host and the *E. coli* HB101 strain which harbored pRK2013 was used as the helper strain for tri-parental mating. Lysogeny broth (LB) liquid medium (tryptone 10 g L^-1^, yeast extract 5 g L^-1^, and NaCl 5 g L^-1^) and solid medium with 1.5% agar was used for conventional cultivation of *E. coli* strains at 37℃, and *P. putida* KT2440 at 30℃. Cetrimide agar was used for selection of *Pseudomonas* strains with a successfully integrated co-integrate after tri-parental mating. When required, 50 mg L^-1^ of kanamycin or 25 mg L^-1^ of gentamycin were added to the medium. All recombinant plasmids in this study were constructed with NEBuilder Hifi DNA Assembly Master Mix (New England Biolabs, Ipswich, Massachusetts, USA). All oligonucleotides were ordered and all sequencing works were done at Eurofins Genomics (Ebersberg, Germany). DNA fragments for plasmid construction were amplified using Q5 DNA polymerase and colony PCRs were performed with One*Taq* 2X Master Mix (New England BioLabs, Ipswich, Massachusetts, USA).

**Table 1.** Plasmids used in this study.

| Plasmid | Description | Source |
| --- | --- | --- |
| pEMG | Suicide vector used for deletions in Gram-negative bacteria; *oriT*, *traJ*, *lacZa*, *oriV(R6K)*, two I-SceI sites; Km<sup>R</sup> | (Martínez-García and de Lorenzo, 2011) |
| pRK2013 | Helper plasmid used for conjugation; *oriV(ColE1)*, *mob(RK2)*, *tra(RK2)*; Km<sup>R</sup> | (Kessler, *et al.*, 1992) |
| pSW-2 | Gm<sup>R</sup>, *oriRK2*, *xylS*, *P<sub>m</sub>→I-SceI* (transcriptional fusion of *I-SceI* to *P<sub>m</sub>*) | (Martínez-García and de Lorenzo, 2011) |
| pSW-2-*sacB* | Derivative of vector pSW-2 carrying *sacB* gene; Gm<sup>R</sup> | In this study |
| pEMG-RIS | Derivative of vector pEMG carrying *P<sub>rhaB</sub>* controlled *I-SceI* and *sacB* gene; Km<sup>R</sup> | In this study |
| pEMG-AIS | Derivative of vector pEMG carrying *P<sub>BAD</sub>* controlled *I-SceI* and *sacB* gene; Km<sup>R</sup> | In this study |
| pEMG-GRIS | Derivative of vector pEMG-RIS, Km<sup>R</sup> is replaced by Gm<sup>R</sup> | In this study |
| pEMG-*pta* | Derivative of vector pEMG carrying TSs for deletion of *pta* (PP_0774) in *P. putida* KT2440; Km<sup>R</sup> | In this study |
| pEMG-RIS-*pta* | Derivative of vector pEMG-RIS carrying TSs for deletion of *pta* (PP_0774) in *P. putida* KT2440; Km<sup>R</sup> | In this study |
| pEMG-RIS-PP_3073 | Derivative of vector pEMG-RIS carrying TSs for deletion of PP_3073 in *P. putida* KT2440; Km<sup>R</sup> | In this study |
| pBG42 | Km<sup>R</sup>, Gm<sup>R</sup>, *oriR6K*, Tn7L and Tn7R extremes, *P<sub>14g</sub>(BCD2)*–*msfGFP* fusion | (Zobel, *et al.*, 2015) |
| pEMG-RIS-0340-*msfGFP* | Derivative of vector pEMG-RIS, for insertion of *P<sub>14g</sub>(BCD2)*–*msfGFP* into the site behind PP_0340; Km<sup>R</sup> | In this study |
| pEMG-RIS-0340-AHHA | Derivative of vector pEMG-RIS, for insertion of a 15.7 kb fragment into the site behind PP_0340; Km<sup>R</sup> | In this study |

**Table 2.** Bacterial strains used in this study.

| Strain | Description | Source |
| --- | --- | --- |
| *Escherichia coli* |  |  |
| DH5α λpir | Cloning host, DH5a strain with λpir lysogen | (Platt, *et al.*, 2000) |
| HB101 | Helper strain, carrying pRK2013 plasmid used for conjugation | (Sambrook, *et al.*, 1989) |
| *Pseudomonas putida* |  |  |
| KT2440 | Wildtype strain, *P. putida* mt-2 derivative cured of the TOL plasmid pWW0 | (Bagdasarian, *et al.*, 1981) |
| KT2440 Δpta | KT2440 with the deletion of *pta* (PP_0774) gene | In this study |
| KT2440 ΔPP_3073 | KT2440 with the deletion of PP_3073 gene | In this study |
| KT2440 0340::GFP | KT2440 inserted with *P*<sub>14g</sub>(*BCD2*)-*msfGFP* at the PP_0340 site | In this study |
| KT2440 0340::AHHA | KT2440 inserted with a 15.7 kb fragment at the PP_0340 site | In this study |

### Plasmid constructions

For the construction of pEMG-RIS and pEMG-AIS plasmids, pEMG vector linearized by PCR amplification or digestion, *sacB* gene with the native promoter and terminator amplified from pLO3 (Lenz and Friedrich, 1998), *P_BAD_* with *araC* gene amplified from pREDCas9 (Li, *et al*., 2015) or *P_rhaB_* with *rhaRS* genes amplified from pEcCas (Li, *et al*., 2021), and the I-SceI gene amplified from pSW-2 (Martínez-García and de Lorenzo, 2011) were assembled by Gibson Assembly. Regarding the sequence in front of the I-SceI gene start codon, it was synthesized on the I*-*SceI gene amplification primer (I-SceI_F) containing the strong RBS (AGGAGG) and 8 bp (AATATACC) between RBS and start codon (Calero, *et al*., 2016). For the construction of pSW-2-*sacB* plasmid, *sacB* gene amplified from pLO3 and pSW-2 vector linearized by PCR were assembled.

As the one plasmid system is based on the pEMG vector, the plasmid construction can still follow the instructions for pEMG vector to achieve the desired editing functions (Martínez-García and de Lorenzo, 2011, Wirth, *et al*., 2020). First of all, pEMG-RIS or pEMG-AIS linearized by PCR amplification (pEMG_V2_F and pEMG_V2_R, Table S1) was used as backbone. Other fragments required in subsequent plasmid assembly are contingent upon the specific objective: 1) for gene deletions, the upstream and downstream targeting sequences TS1 and TS2 (∼500 bp of each) of the deletion area amplified from *P. putida* KT2440 genome, are required; 2) for gene insertions, additional DNA fragments are needed and should be positioned between TS1 and TS2; 3) for point mutations and short insertions, the mutation can be introduced within the primer overhangs. For more details on the primer design with pEMG vector, please refer to the provided descriptions by (Wirth, *et al*., 2020).

### Tri-parental mating and cetrimide agar selection

For plasmid delivery, the conjugation (tri-parental mating) option is highly recommended as the method of choice (Aparicio, *et al*., 2019). Briefly, this process includes overnight cultivation of the plasmid donor strain (*E. coli* DH5a λ*pir* harboring the editing plasmid), helper strain (*E. coli* HB101 harboring pRK2013) and the acceptor strain (*P. putida* KT2440 and its derivatives) in 4 mL LB medium with corresponding antibiotics. Then, 200 μL of each culture was mixed in a 1.5 mL reaction tube and washed once with LB medium after centrifugation. Finally, it was resuspended with 200 μL LB medium and 20 μL of the mixture was plated by sterile inoculation loop on the pre-dried LB plate (Wirth, *et al*., 2020) at 30℃ for 6∼8 h. For co-integrated strain selection, biomass on plate agar surface was taken by a pipette tip and resuspended in 600 μL of LB medium. Next, 100 μL of that suspension was spread on a cetrimide agar plate supplemented with 50 mg/L of kanamycin and then incubated at 30°C for 14∼16 h. Colony PCR was used for the verification of co-integration.

### Induction of I-SceI endonuclease and selection of plasmid-free mutants

To introduce the I-SceI mediated double-strand breaks, colonies of *P. putida* KT2440 strain co-integrated with pEMG-RIS/pEMG-AIS vector from cetrimide agar plate mentioned above were picked into 4 mL LB medium supplemented with 10 mM rhamnose (pEMG-RIS) or 100 mM arabinose (pEMG-AIS) in test tubes. After 4∼12 h, that culture was diluted and spread on the LB plates supplemented with 10% sucrose. Then they were incubated overnight and several colonies were picked for PCR verification. Further verification was done by PCR amplification of the related sequences using Q5 DNA Polymerase and was sent for sequencing after gel purification.

## Results and Discussion

### Design a user-friendly genome editing tool for Pseudomonads

The pEMG/pSW system, developed by Martínez-García et al. (2015), has been extensively employed in gram-negative bacteria. It functions as a highly reliable and standardized tool that has greatly facilitated research involving Pseudomonads (Martínez-García, *et al*., 2014, Wynands, *et al*., 2018). To further expedite the genome editing process, it’s valuable to analyze the optimizable operations of the existing system and contemplate the stepwise optimization. As shown in Fig. 1, it’s evident that the curing of the pSW-2 plasmid constitutes the most time-intensive aspect of this original system. Generally, a minimum of 6 successive passages (Wirth, *et al*., 2020) in LB medium without selection pressure is required, which takes at least 2∼3 days. Another seemingly laborious part is the delivery of the second plasmid pSW-2, either using conjugation or electroporation.

**Fig. 1.**
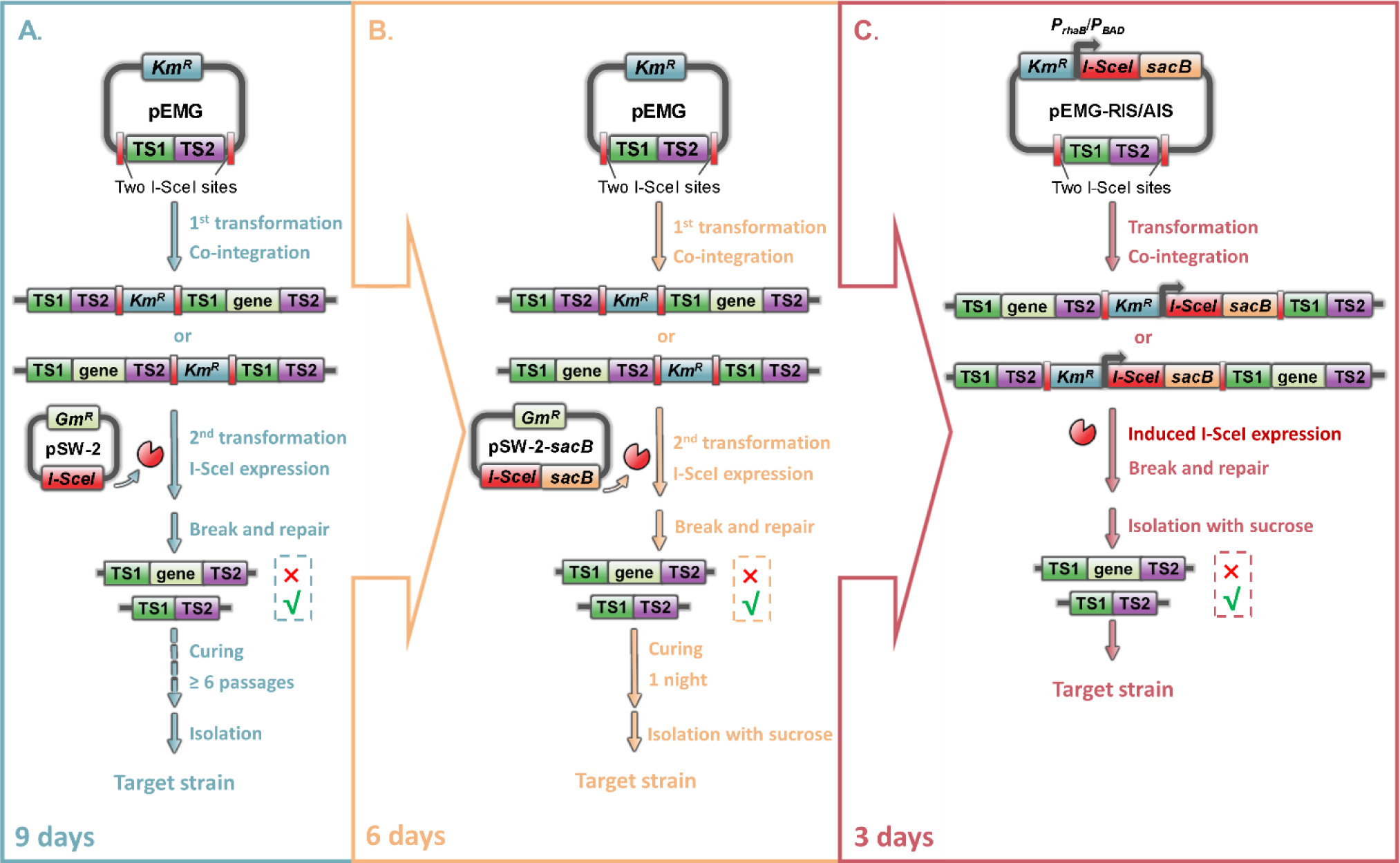
Overview of the workflow with the original pEMG/pSW system and the implementation of two alternative strategies. (A) The original pEMG plasmid bearing a Km^R^ marker, an origin of replication *oriR6K*, an *oriT*, and two I-SceI sites on the backbone. For gene deletions, the upstream and downstream targeting sequences (TS1 and TS2) are assembled into this plasmid between the two I-SceI sites. Firstly, this suicide vector was co-integrated into *P. putida* KT2440 genome via tri-parental mating. Secondly, the I-SceI endonuclease was introduced by the delivery of the pSW-2. Then, the resulting DSB caused by I-SceI were repaired by homologous recombination. Successful co-integrate release it tested by kanamycin sensitivity. Next, after the genotype verification through colony PCR, a plasmid curing procedure was conducted. (B) Base on the pEMG/pSW system, a counterselection marker *sacB* was introduced onto pSW-2, resulting in pSW-2-*sacB*. Then, the utilization of sucrose facilitated the isolation of plasmid-free strains, avoiding the curing of pSW-2 and reducing the overall time required for the procedure from 9 to 6 days. (C) In a one plasmid system, both the I*-*SceI gene controlled by *P_rhaB_* or *P_BAD_* and *sacB* gene were assembled into the backbone of the pEMG vector, named pEMG-RIS and pEMG-AIS, respectively. Then, after co-integration, I-SceI was expressed with the direct induction of rhamnose or arabinose in LB liquid cultures and plated on LB agar plates supplemented with 10% sucrose. The streamlined procedure was accomplished within 3 days.

### Accelerating the plasmid curing process of pSW-2

To accelerate the process of plasmid curing, the *sacB* gene, which is commonly employed as a counterselection marker, has been considered and incorporated into the pSW-2 plasmid. The resulting plasmid is named pSW-2-*sacB* (Fig. 2A). Thus, the utilization of sucrose during the plasmid curing procedure enables the isolation of plasmid-free mutants. To validate the optimized plasmid, we conducted an experiment involving the deletion of the *pta* gene (PP_0774, 2.1 kb, encoding the phosphate acetyltransferase) in *P. putida* KT2440. For the deletion of *pta* gene, it worked as well as the original pSW-2 plasmid. To evaluate the plasmid curing efficiency, a dilution series of a liquid culture of a correct colony was spread on LB agar plates supplemented with different concentrations of sucrose (0%, 0.5%, 1%, 2%). As shown in Fig. S1A, many colonies grew on the plate lacking sucrose, whereas significantly fewer colonies grew on the plates with sucrose. Subsequently, colonies were randomly picked from the plates and tested for the loss of pSW-2-*sacB* using LB-Gm plates. As expected, all colonies from the plates supplemented with sucrose exhibited plasmid loss (Fig. S1B). These results indicated that the pSW-2-*sacB* plasmid not only can facilitate the isolation of plasmid-free strain on LB-sucrose plates but also works as good as the pSW-2. This obviates the need of multiple passages of strain cultivation procedures, resulting in a time-saving of approximately three days.

**Fig. 2.**
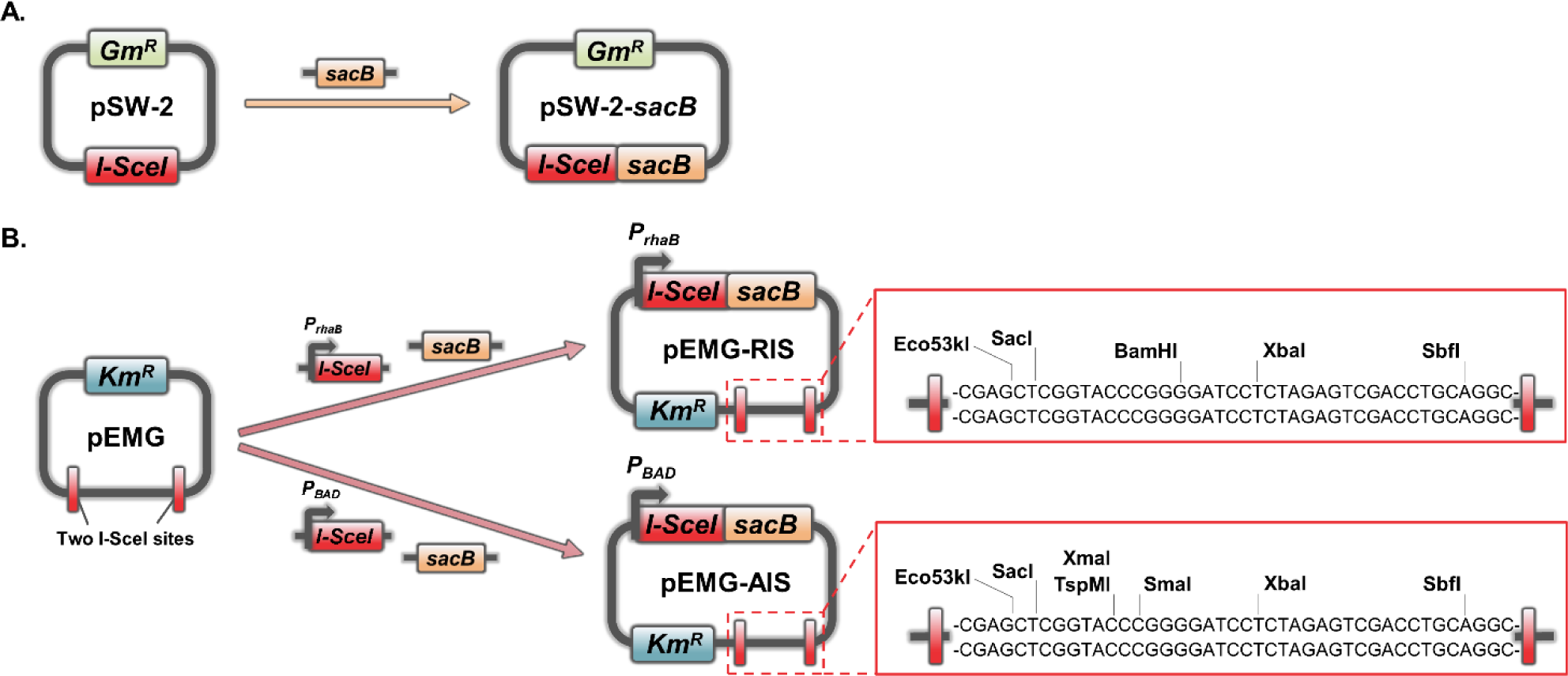
Design of the construction of pSW-2-*sacB* (A), as well as the construction of pEMG-RIS and pEMG-AIS (B). The DNA sequences flanked by I-SceI sites depict the multiple cloning sites in both plasmids. These sites enable plasmid linearization of pEMG-RIS or pEMG-AIS by restriction endonuclease digestion.

### Constructing a one plasmid system

Creating a one plasmid system was then considered to further streamline the process. However, the system requires incorporating the I-SceI gene and the two I-SceI sites in a single vector, which may result in “self-cleavage”. To avoid this issue, it’s crucial to ensure strict control of the I-SceI expression. Calero *et al*. (2016) characterized several inducible promoter systems in *P. putida* and demonstrated that the xylS/*P_m_* promoter exhibits a high leakage. Whereas the promoters *P_rhaB_* and *P_BAD_* show a very low level of basal expression and are therefore suitable for inducible expression of heterologous genes (Calero, *et al*., 2016). For this reason, we replaced the xylS/*P_m_* promoter, used for I-SceI expression on plasmid pSW-2, by AraC/*P_BAD_* or RhaRS/*P_rhaB_*. The inducible promoters were fused with I-SceI and integrate it into the pEMG vector alongside the *sacB* gene, yielding pEMG-RIS or pEMG-AIS (Fig. 2B). This allowed us to strictly control I-SceI expression by the addition of rhamnose (pEMG-RIS) or arabinose (pEMG-AIS). Moreover, unique restriction sites remain in the multiple cloning site (MCS) between the I-SceI sites, enabling plasmid linearization by restriction endonuclease digestion (Fig. 2B). Consequently, a one plasmid system has been established, which theoretically enables one round of genome editing process within three days (Fig. 1C and Fig. 4A).

To verify the applicability and efficiency of the newly constructed plasmids, the gene *pta* in *P. putida* KT2440 should be deleted. The obtained plasmids pEMG-RIS-*pta* and pEMG-AIS-*pta* were integrated into the genome and afterwards I-*Sce*I expression induced by either rhamnose or arabinose to introduce DSB in the co-integrate. Both plasmids showed satisfactory performance, achieving positive rate of 21/28 and 12/28, respectively. Notably, the plasmid containing *P_rhaB_* performed better compared to the *P_BAD_* inducible variant, as confirmed through repeated experimentation. Calero *et al*. showed that *P_rhaB_* promoter in *P. putida* KT2440 is stronger and require lower concentration of inducer (Calero, *et al*., 2016), which may contribute to the better performance. Consequently, the pEMG-RIS vector was selected for subsequent demonstrations.

However, upon integration into the genome, the functional efficiency of the SacB and I-*Sce*I can be down-regulated. In contrast to the consistently high expression on plasmids, expression on the genome is not only down-regulated due to single copy, but also influenced by the genomic context (Espah Borujeni, *et al*., 2014, Englaender, *et al*., 2017, Köbbing, *et al*. submitted). We have observed some co-integrated strains on the LB plate supplemented with 1% sucrose, which revealed that some cells didn’t demonstrate targeted cleavage mediated by I-SceI after induction when utilizing the one plasmid system. Moreover, the selection condition of 1% sucrose employed for pSW-2-*sacB* was inadequate here, which led to a re-evaluation of counterselection on LB plate supplemented with different concentrations of sucrose (see Fig. S2). As expected, the growth of uninduced strains decreased substantially with increasing sucrose concentrations and a 10% sucrose concentration sufficiently fulfills the counterselection requirements of the genome editing system in *P. putida* KT2440.

### Targeted integration of msfGFP in the genome of P. putida KT2440

To visualize the efficiency and reliability of our one plasmid system, we integrated a gene expression cassette containing *msfGFP* in the genome *P. putida* KT2440. For targeted integration, we chose the genomic locus next to PP_0340 (Köbbing, *et al*. submitted). To facilitate the insertion, a pEMG-RIS vector containing the upstream and downstream homologous regions was constructed. The expression cassette consists of a *msfGFP* reporter gene driven by a constitutive σ^70^-dependend promoter *P_14g_* (Zobel, *et al*., 2015), a *BCD2* elements (Mutalik, *et al*., 2013) and *T_0_* terminator (Fig. 3).

**Fig. 3.**
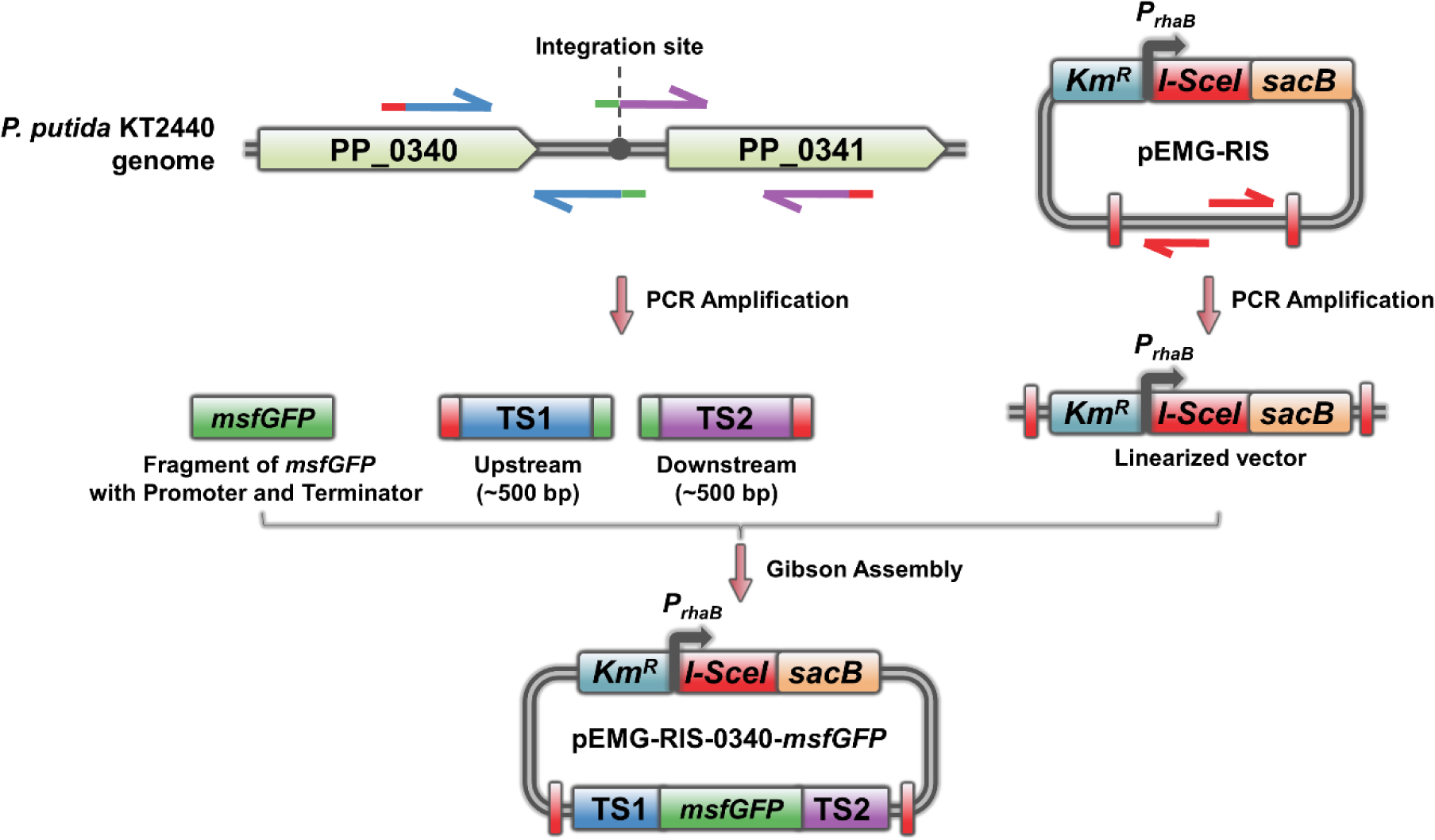
Workflow for constructing the pEMG-RIS-0340-*msfGFP* plasmid for gene insertion in *P. putida* KT2440.The upstream and downstream targeting sequences (TS1 and TS2) amplified by two pairs of primers from *P. putida* KT2440 genome, linearized pEMG-RIS vector, and the *msfGFP* gene with *P_14g_* promoter and *T_0_* terminator were assembled via Gibson Assembly. Then, the assembled pEMG-RIS-0340-*msfGFP* plasmid is introduced into *E. coli* DH5a λ*pir*.

The resulting *E. coli* DH5a λ*pir* strain containing plasmid pEMG-RIS-0340-*msfGFP* was then used as donor strain as described in Fig. 4A. After 4 h induction, 50 μL of that culture was spread on LB plates with or without 10% sucrose selection and incubated overnight, respectively. Then, photos were taken on a blue tray (Fig. 4B), colonies counted and verified by colony PCR. Through PCR, we confirmed that only the co-integrate released strains can grow in the presence of 10% sucrose. While colonies derived from normal LB plate resulted in a mixture of co-integrate strain, integrate or wildtype (Fig. 4B). The evaluation of integration efficiency of the expression cassette alone into the genome was then performed through direct observation of clones with or without green fluorescence on a blue tray (Fig. 4B). On plates supplemented with 10% sucrose, 50.6 ± 0.2% (140/277) of the colonies (green spots) contained the desired *msfGFP* genomic integration. And on plates without sucrose, only 4.9 ± 0.1% (137/2816) of the colonies (dark spots) were the wildtype strains which means most of the colonies were co-integrate unreleased. These results showed that the addition of 10% sucrose to LB plates effectively functioned as a counterselection, resulting in a notable decrease in the number of PCR samples needed to obtain the target strain. As expected, these results provided a comprehensive and visual validation of our method, achieving the anticipated theoretical efficiency of 50%.

**Fig. 4.**
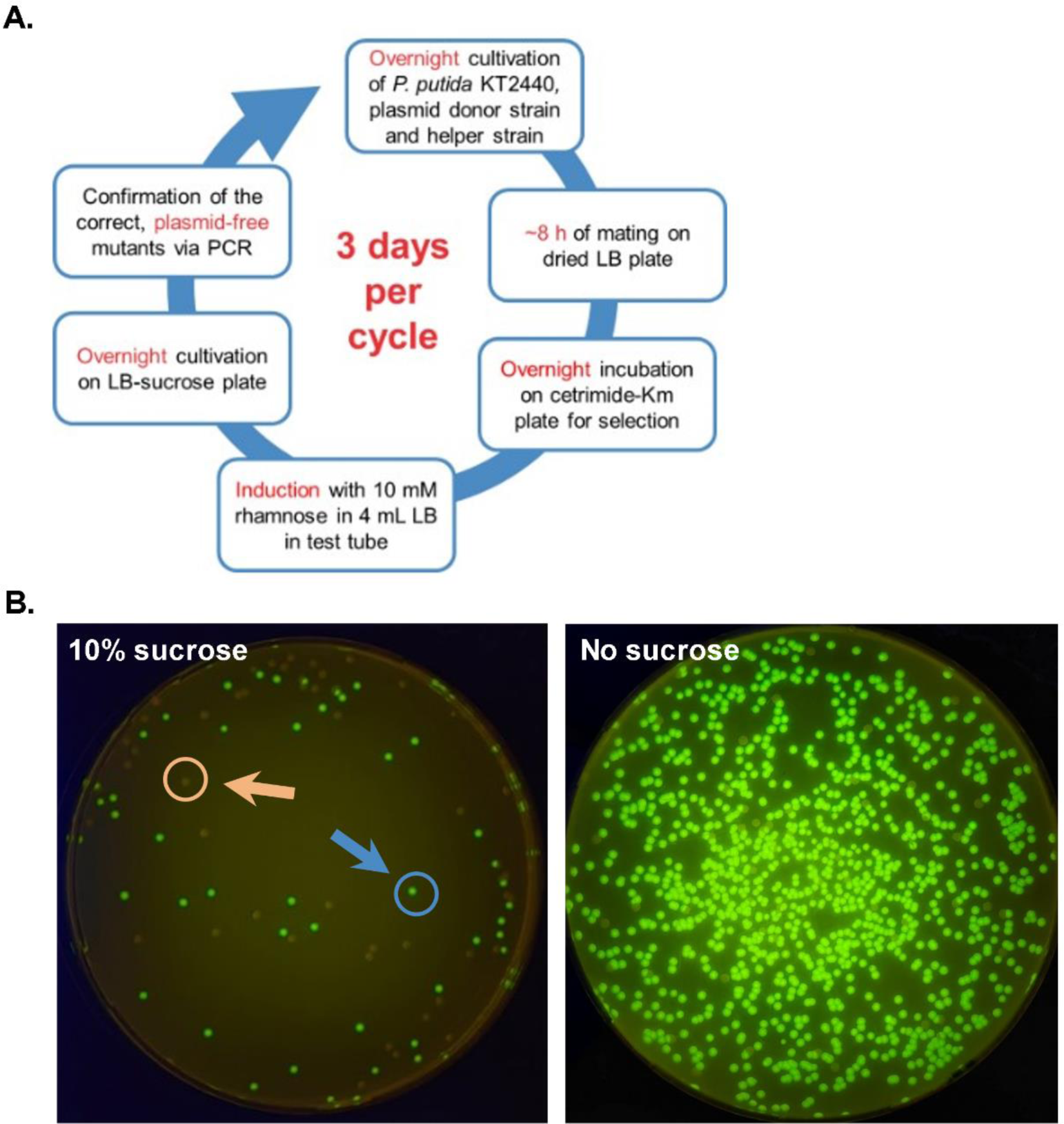
A 3-day workflow (A) for the integration of the *msfGFP* gene into *P. putida* KT2440 using the one plasmid system. The plasmid donor strain is *E. coli* DH5a λ*pir* harboring pEMG-RIS-0340-*msfGFP* and the helper strain is *E. coli* HB101 harboring pRK2013. (B) The plates demonstration of *msfGFP* integration with (left, on blue tray) or without (right, on blue tray) sucrose selection. On the left plate, only *msfGFP* gene inserted (green spots) and wildtype (dark spots) *P. putida* KT2440 strains grown. On the right plate, green spots indicate colonies of *msfGFP* gene inserted or co-integrated strains whereas the dark spots are wildtype *P. putida* KT2440 strains.

### Insertion of a large fragment using the one plasmid system

Strain engineering often requires the integration of larger fragments into the genome, which can be challenging for genome editing methods. To evaluate the efficiency of the one plasmid system in addressing this challenge, we conducted an experiment involving the insertion of a 15.7 kb fragment into the genome next to PP_0340 (Köbbing, *et al*. submitted). Surprisingly, despite the considerable size of the resulting integration plasmid (24.8 kb), it showed a commendable performance with a positive rate of 10/96. Other colonies are wildtype, seemingly suggesting that homologous recombination repair here prefers to generate a shorter genome. Consequently, we can confidently assert that this one plasmid system is fully capable of meeting any conventional gene insertion requirements.

### The location of gene in the genome can influence its modification

We have observed that the effectiveness of the SacB and I-SceI in the one plasmid system can be influenced by the genomic context. The system was not working perfectly in some cases compared to the gene insertion work at PP_0340 site, which was defined as a “hot spot” in the genome (Köbbing, *et al*. submitted). We evaluated the co-integrate release rate of the deletion of a “cold site”, PP_3073, compared with the deletion of *pta* gene and gene insertion at the PP_0340 site. Our findings indicate that longer induction times led to increasingly higher release rates of each co-integrate. The release rates at the PP_3073 site was significantly lower compare to PP_0340 site and *pta* gene (Fig. 5). Furthermore, the strain with the co-integrate at the PP_3073 site could still grow on LB plates supplemented with 10% sucrose. However, the size of colonies was much smaller than those colonies with co-integrate release (Fig. S3). These results indicated that the genome locus can significantly affect genome editing with the one plasmid system, and an induction time of 12 h or even longer might be more effective.

**Fig. 5.**
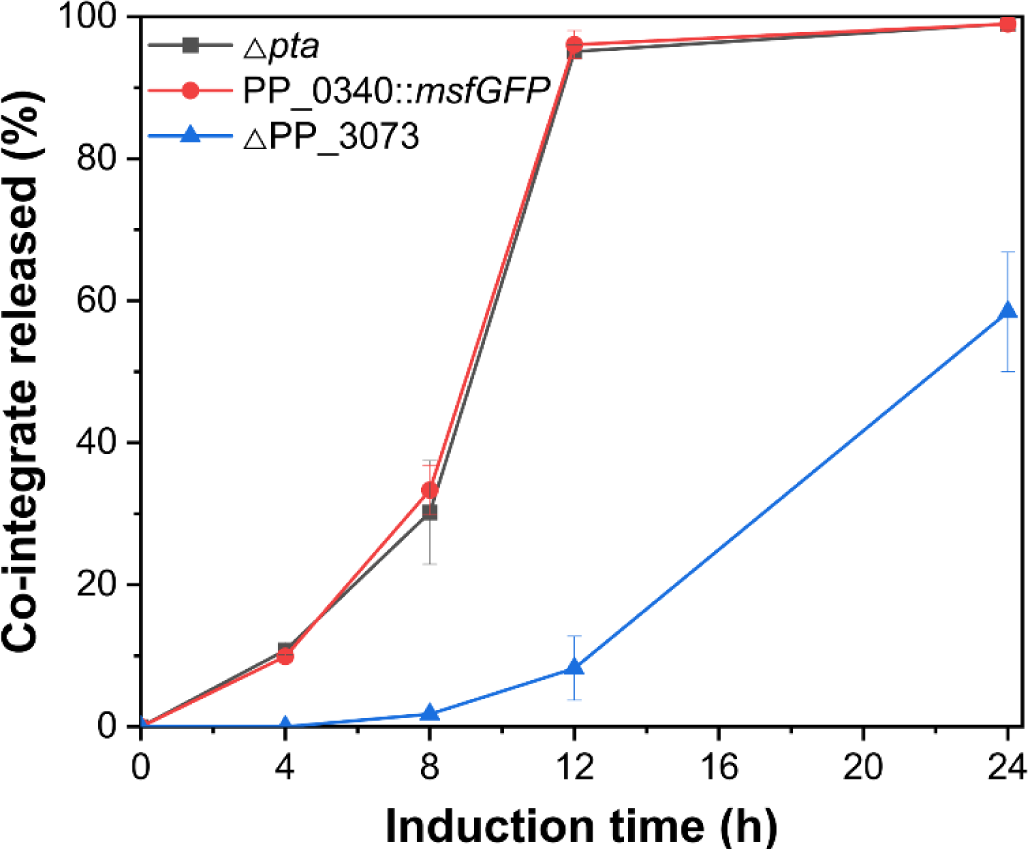
Time dependency of co-integrate release. Different times of I-SceI induction during gene insertion at PP_0340, or gene deletion of *pta* and PP_3073 were tested. The experiment was conducted by cultivating the co-integrated strains in LB medium supplemented with 10 mM rhamnose. These samples were then diluted and spread on LB plates. After overnight incubation, thousands of single colonies were picked and tested by LB-Km plates. The colonies that were unable to grow on LB-Km plates correspond to cells that released the co-integrate.

## Conclusion

In this study, we demonstrate a one plasmid genome editing tool, which accelerates the engineering cycle from nine down to three days. Based on the pEMG backbone, a tightly controllable I-SceI gene and a constitutive *sacB* gene were assembled on a single plasmid. Consequently, the requirement of a second plasmid transformation and a laborious plasmid curing process can be circumvented. Notably, the new pEMG-RIS single plasmid system is also reliable for large fragment insertion.

In more detail, the inclusion of a counterselection marker is of significant importance. The SacB as an additional counterselection marker demonstrated remarkable efficiency for gene deletion and insertion. In the literature, SacB performance was questioned in some applications (Galvão and de Lorenzo, 2005, Luo, *et al*., 2016). Hence, to broaden the applicability of the pEMG-RIS system, one might explore alternative counterselection markers, such as *pheS* (Miyazaki, 2015).

To further accelerate genome editing, we attempted to simultaneously execute two gene deletions by employing pEMG-RIS plasmids containing distinct antibiotic markers. Regrettably, this attempt proved unsuccessful, potentially due to the long identical sequences on the plasmid backbone, which may impede the co-integration.

To effectively address the growing demand for genome editing in *P. putida*, we present here our novel one plasmid system pEMG-RIS. We hope that this 3-day method can expedite the exploration of *P. putida* and expand its use in biological and environmental applications. The pEMG-RIS will be available via Addgene.

## Author Contributions

H.M. designed and conducted the experiments, and wrote the manuscript; S.K. and L.M.B. supported the experimental design, revised the manuscript and supervised the study. L.M.B. provided the funding for this work. All authors have approved the final version of the manuscript.

## Acknowledgments

This work was financially supported by the China Scholarship Council (CSC), and the European Union’s Horizon 2020 research and innovation program under grant agreement no. 870294 (MIX-UP project).

## ORCID

*Hao Meng,* https://orcid.org/0009-0003-4649-2047

*Sebastian Köbbing,* https://orcid.org/0000-0002-4606-4961

*Lars M. Blank* https://orcid.org/0000-0003-0961-4976

## Supporting Information

The Supporting Information for this article can be found online in the Supporting Information section at the end of this article.

