## Supporting Information for "Establishing a straightforward I-SceI mediated recombination one plasmid system for efficient genome editing in *P. putida* KT2440"

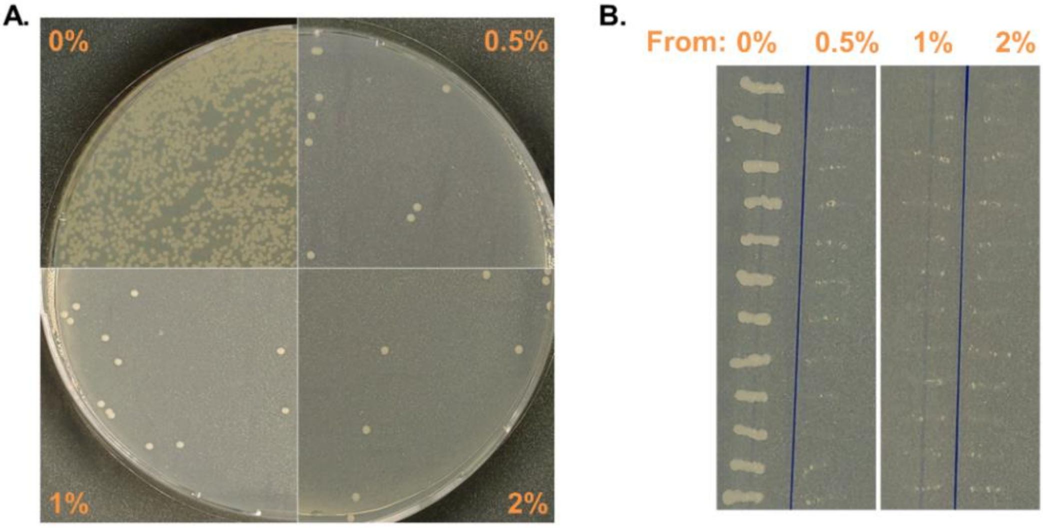


Fig. S1 Evaluation of SacB as counterselection marker on pSW-2 plasmid. After the cultivation of one correct colony containing the pSW-2-*sacB* plasmid in LB medium for 12 h, the culture was diluted to $OD_{600}$ 0.001 and spread on LB plates (A) supplemented with four different concentrations of sucrose (0%, 0.5%, 1%, 2%). To verify the selection, colonies from the four plates are randomly selected and tested on LB-Gm plates (B).

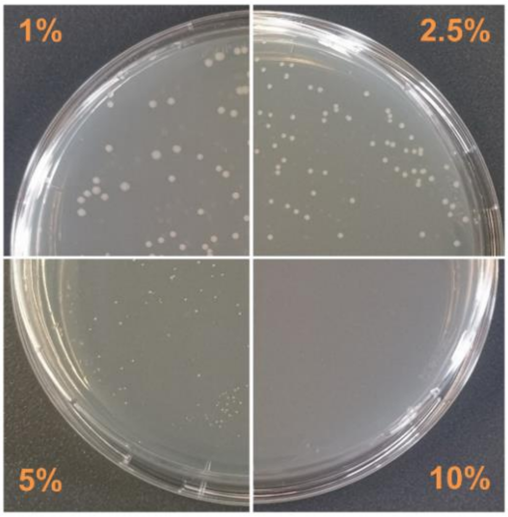


Fig. S2 Evaluation of SacB as a counterselection marker after co-integrated on the genome of *P. putida* KT2440. Overnight cultivation of the *P. putida* KT2440 strain with a co-integrated pEMG-RIS-*pta* in LB medium without the addition of inducer. Subsequently, the uninduced fresh culture was diluted to $OD_{600}$ 0.001 and spread on LB plates supplemented with four different concentrations of sucrose (1%, 2.5%, 5%, 10%), and then incubated at 30℃ for 24 h.

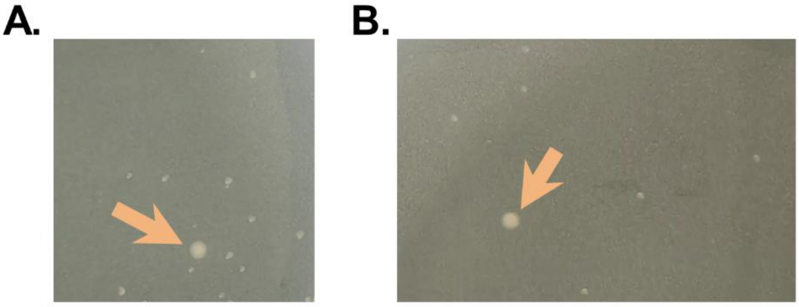


Fig. S3 Counterselection results during the genomic modification at the "cold spot", PP_3073. After being induced with 10 mM rhamnose for 8 h (A) or 12 h (B), the cell cultures were diluted to $OD_{600}$ 0.001and spread onto the LB plates supplemented with 10% sucrose. The co-integrated strain can still grow in the presence of 10% sucrose, but formed much smaller colonies compared to colonies with successfully released co-integrates (indicate by orange arrows).

Table S1 Oligonucleotides used in this study

| Primers | Sequence (5′→3′) | Source |
|---|---|---|
| pSW-insert-F | GTGATAATCACTCGCACGCTGG | This study |
| pSW-insert-R | GAACTGGCCAGCAAGGTCAG | This study |
| pLO3-sacB-F | GACCTTGCTGGCCAGTTCCTTGCGGAGAACTGTGAATGC | This study |
| pLO3-sacB-R | AGCGTGCGAGTGATTATCACCGCTACGATCCTTTTTAACCCAT CAC | This study |
| sacB-seq-F | CAGCATATCATGGCGTGTAATATGG | This study |
| sacB-seq-R | GACAAACAGAGGATTCTACGCAGAC | This study |

| | | |
|---|---|---|
| pEMG-V2-F | AGTCGACCTGCAGGCATG | This study |
| pEMG-V2-R | CTAGAGGATCCCCGGGTACC | This study |
| 0340-TS1-F | GGTACCCGGGGATCCTCTAGCGCGACCTGGAAAAGCTG | This study |
| 0340-TS1-GFP-R | CTTGTCAATGGGCTTAATTAAAGCACTGCTCAGGGCCTGTG | This study |
| p14g-GFP-F | CTTTAATTAAGCCCATTGACAAGGCTC | This study |
| GFP-term-R | ATCTGACGTCCTTGGACTCCTG | This study |
| 0340-TS2-GFP-F | GGAGTCCAAGGACGTCAGATTGCAATCCCTGTGGGAGC | This study |
| 0340-TS2-R | TGCATGCCTGCAGGTCGACTTTGCGCACATCGTTCAGCAG | This study |
| pEMG-seq-F | CATTCAGGCTGCGCAACTG | This study |
| pEMG-seq-R | GTGAGTTAGCTCACTCATTAGGCAC | This study |
| pta-seq-F | GCTATTACGCCGACAAGCAG | This study |
| pta-seq-R | GTACTCGCCGTAGGCCTC | This study |
| pta-TS1-F | GGTACCCGGGGATCCTCTAGCGCACAGATGGCCAACAC | This study |
| pta-TS1-R | TAAAAGCAGAGGCTTCTTTGCAAGC | This study |
| pta-TS2-F | CAAAGAAGCCTCTGCTTTTACATGCGTGCTTCTCGAAATAAGC | This study |
| pta-TS2-R | TGCATGCCTGCAGGTCGACTCATTCAGCTCGTCACCAGC | This study |
| pEMG-V-araBAD | CTTCGCCAACTATTGCGCTGATCTGGACAAGGGAAAACGCAA G | This study |
| pEMG-V-rhaB | GATACAAGAGCCATAAGAACCATCTGGACAAGGGAAAACGCA AG | This study |
| ParaBAD-F | CAGCGCAATAGTTGGCGAAG | This study |
| ParaBAD-R | GGTATATTCCTCCTATCGCCTCAGCATCCAAAAAAACGGGTAT GGAGAAACAGTAGAGAGTTGCG | This study |
| PrhaB-F | GGTTCTTATGGCTCTTGTATCTATCAGTGAAG | This study |
| PrhaB-R | GGTATATTCCTCCTATCGCCTCAGCGAATTTCATTACGACCAGT CTAAAAAGC | This study |
| I-SceI-F | GGCGATAGGAGGAATATACCATGAAAAACATCAAAAAAAACC AGGTAATGAACC | This study |
| I-SceI-RW | TCGACTTATTATTTCAGGAAAGTTTCGGAGG | This study |
| sacB-F | CTTTCCTGAAATAATAAGTCGACTTGCGGAGAACTGTGAATGC | This study |
| sacB-R | CGCTACGATCCTTTTTAACCCATCAC | This study |
| pEMG-V-sacB | GGTTAAAAAGGATCGTAGCGAGCCCAGTAGCTGACATTCATC | This study |
| I-SceI-seq-1 | CATCACGAGAACGGATGTAAGCATC | This study |
| I-SceI-seq-2 | GTCTGCGTAACAAATTCCAACTGAAC | This study |
| pEMG-seq-3 | GACATGGGAATTAGCTTCACGC | This study |
| pEMG-seq-4 | CCAGTTTACTTTGCAGGGCTTC | This study |
| RP-Gm-Fragment-F | GGGGTGGGCGAAGAACTTCTAGGGCGGCGGATTTG | This study |

| | | |
|---|---|---|
| RP-Gm-Fragment-R | GTAGCTTGCAGTGGGCTTACCTTGACATAAGCCTGTTCGGTTC | This study |
| RP-Km-pEMG-V-R | AGTTCTTCGCCCACCCC | This study |
| RP-Km-pEMG-V-F | GTAAGCCCACTGCAAGCTACC | This study |
| HM-seq-4 | GCCTGATAGCCGTTGATCTTGG | This study |
| HM-seq-6 | CGAGATTGGGCTGATCGATG | This study |
| PP0340-int-seq-F | CTTCGAACGCTACCAGCAGAAC | This study |
| PP0340-int-seq-R | CAGTACCGGACGGTCCAG | This study |
| 0340-int-TS1-R-AHHA | GACTACACTTAAGTGAAGCTCACTGCTCAGGGCCTGTG | This study |
| 0340-int-TS2-F-AHHA | CGGCGTTTCAGGTCAAGCTATAAGAATTCGAGCTCGGTACCC | This study |
| PP_3073-seq-F | GGTAGCGCACTTCGACATC | This study |
| PP_3073-seq-R | TGGCCGAAGGCACCAAAAG | This study |
| PP_3073-TS1-F | GGTACCCGGGGATCCTCTAGGGCACATACACCACCACCG | This study |
| PP_3073-TS1-R | TGAAGGCAAAACTGCACTCGATGGTGGCTGGTTGG | This study |
| PP_3073-TS2-F | GAGTGCAGTTTTGCCTTCAAGG | This study |
| PP_3073-TS2-R | TGCATGCCTGCAGGTCGACTCTTGCTGGCAGCCATGAAC | This study |
